# Pan-Cancer Genetic Profiles of Mitotic DNA Integrity Checkpoint Protein Kinases

**DOI:** 10.1101/2024.11.15.623822

**Authors:** Ayana Meegol Rasteh, Hengrui Liu, Panpan Wang

**Author notes:** These authors contribute equally (Ayana Meegol Rasteh, Hengrui Liu). Corresponding: Hengrui Liu Panpan Wang.

## Abstract

**BACKGROUND:** The mitotic DNA integrity checkpoint signaling pathway is potentially involved in cancers that regulate genomic stability where protein kinases play a pivotal role. 16 total protein kinase genes are involved in this pathway: ATM, BRSK1, CDK1, CDK2, CHEK1, CHEK2, MAP3K20, NEK11, PLK1, PLK2, PLK3, PRKDC, STK33, TAOK1, TAOK2, and TAOK3. This study aims to provide pan-cancer profiles of the protein kinases in mitotic DNA integrity checkpoint signaling gene set for potential prognostic and diagnostic purposes, as well as future potential therapeutic targets for cancer in a clinical setting.

**METHODS:** Multi-omic data was acquired for the 16 genes; over 9000 samples of 33 types of cancer were analyzed to create pan-cancer profiles of SNV, CNV, methylation, mRNA expression, pathway crosstalk, and microRNA regulation networks.

**RESULTS:** The SNV profile showed that most of these genes have a high SNV mutation frequency across some cancer types, such as UCEC and SKCM. The CNVs of some of these genes are associated with the survival of UCEC, KIRP, and LGG. BRCA, KIRC, LUAD, and STAD might be affected by the mRNA expression of these genes which might involve regulation of copy number, methylation, and miRNA. In addition, these genes also cross-talk with some known cancer pathways.

**CONCLUSION:** The protein kinases in mitotic DNA integrity checkpoint signaling may play a role in cancer development and, with adequate research, could potentially be developed as biomarkers for cancer diagnosis and prognosis. However, further efforts are necessary to validate their clinical value for diagnosis and prognosis and to develop practical applications in clinical settings. Nevertheless, these pan-cancer profiles offer a better overall understanding as well as useful information for future reference regarding mitotic DNA integrity checkpoint signaling in cancer.

## INTRODUCTION

Cancer is a complex disease that impacts many worldwide. In the United States alone, it is estimated that approximately 1,958,310 people will be newly diagnosed with cancer in 2023, and 609,820 people will die from cancer [1]. Despite the recent advancements in cancer research, there is still a need for more effective treatment options. Molecular analyses have deemed that traditional categorization of cancer into groups based on the tissue where it occurs can be restrictive [2], as tumors originating from different organs can manifest similar characteristics, whilst tumors from the same organ may exhibit differences. The pan-cancer analysis allows for the comparing and contrasting of characteristics across many cancers[3], offering a comprehensive understanding of them. The Cancer Genome Atlas (TCGA) plays an important role in pan-cancer research, as it provides thorough data for analyzing diverse cancer genomes [4]. TCGA data has been widely used in both single-cancer type studies[5–12] and pan-cancer studies[13–18].

Under typical circumstances, mitosis generally results in the formation of two cells with the correct chromosome count, known as euploidy. However, disrupted mitosis can present itself in an abnormal chromosomal number, aneuploidy, which is a known hallmark of cancer [19]. The mitotic DNA integrity checkpoint in normal cells acts as a method for maintaining genomic stability, as it detects DNA irregularities and prevents the spread of damaged genomes [20]. However, a failure of this process can cause aneuploidy, paving the way for potential tumorigenesis and metastasis [21,22]. Protein kinases play a crucial role in mitotic DNA integrity checkpoint signaling as they can regulate the process directly or trigger signals to other downstream kinases, both of which help maintain the integrity of the DNA and avoid cancerous cells[23].

Based on the Gene Ontology (GO) database, the ancestor chart shows where mitotic DNA integrity checkpoint signaling pathway is located on the GO structure (Fig.1A). A total of 87 genes are included in this pathway and based on their "gene families" categorized by GSEA database (Fig.1B), we obtained 16 total protein kinase genes that are involved in the mitotic DNA integrity checkpoint signaling, including ATM, BRSK1, CDK1, CDK2, CHEK1, CHEK2, MAP3K20, NEK11, PLK1, PLK2, PLK3, PRKDC, STK33, TAOK1, TAOK2, and TAOK3. Some of these genes, including ATM and MAP3K20, are positive regulators in DNA repair within the realm of cancer [24,25]; some are also negative regulators, such as CDK1, and CDK2 [26,27]. CDK2 has been previously reported as a glioma biomarker[9]. Understanding the role of these genes in cancer is crucial for grasping the complexities of cancer growth related to the mitotic DNA integrity checkpoint signaling protein kinases. Developing this pathway as a potential therapeutic target offers a way to improve clinical cancer treatments.

**Figure 1:**
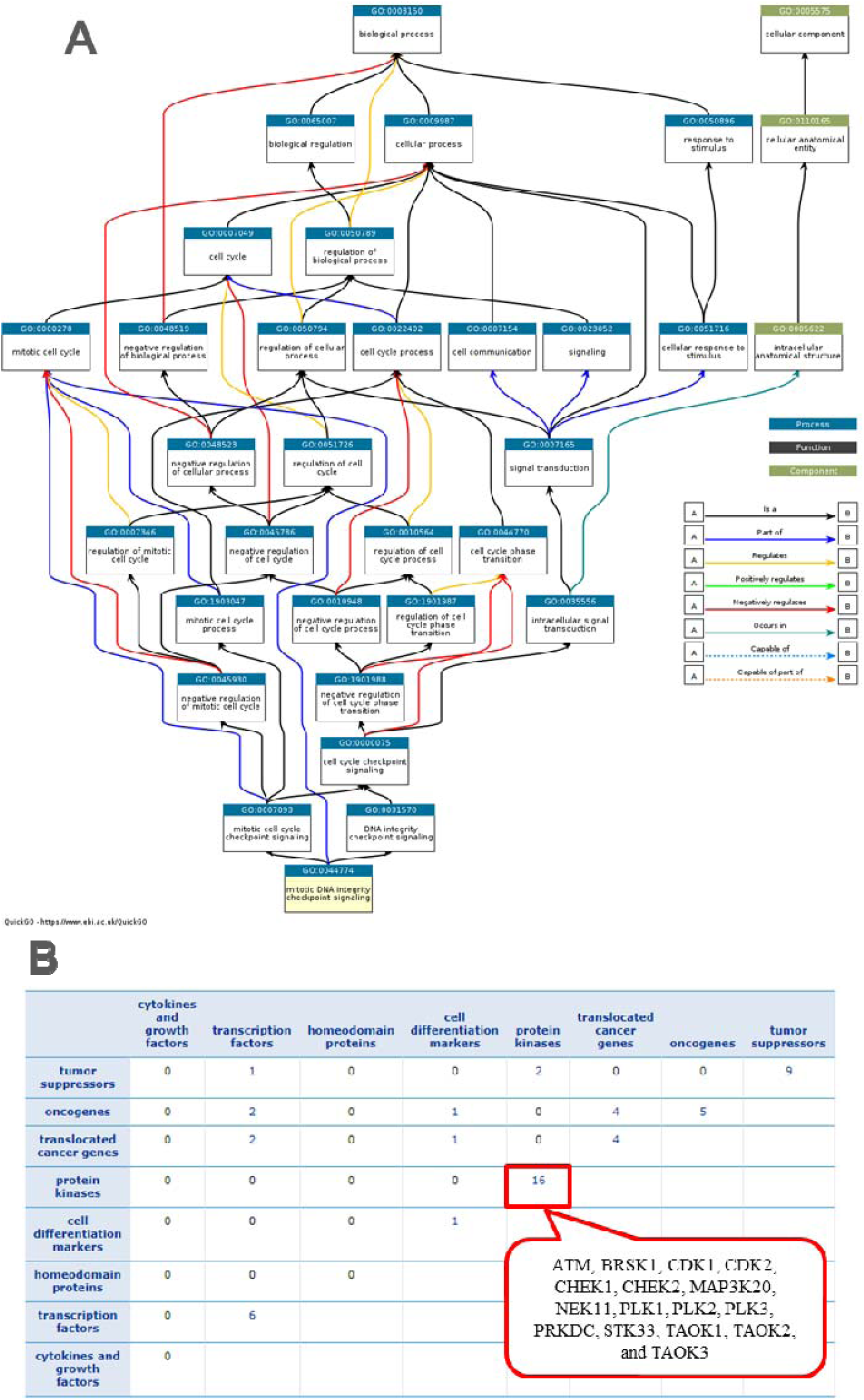
A: Ancestor Chart of mitotic DNA integrity checkpoint signaling pathway from Gene Ontology (GO) database. B: The table providing a functional overview of the gene sets by categorizing their genes into a small number of carefully chosen "gene families". Members of these "gene families" share a common feature such as homology or biochemical activity. They do not necessarily have common origins. The source of each "gene family" definition follows GSEA database.

Utilizing multi-omic profiling data, we categorized these 16 genes across over 9000 samples from 33 different types of cancer. This comprehensive pan-cancer study revealed the diverse expression regulations of the protein kinases within the mitotic DNA integrity checkpoint signaling gene set; it also examined potential associations prevalent cancer signaling pathways, giving an overall view of these genes for future reference. We believe that these profiles will provide genetic information that will be useful for future cancer studies.

## METHODS

### DATA ACQUISITIONS

The data for the single nucleotide variant (SNV), copy number variant (CNV), RSEM normalized mRNA expression data, survival data, and methylation data with clinical information were collected from The Cancer Genome Atlas database (TCGA) [4]. The reverse phase protein array (RPPA) data was gathered from the cancer proteome atlas (TCPA) [28,29]. The regulation data for the miRNA was acquired from databases with experimentally validated scientific papers.

The databases include mir2disease data [30], miRTarBase [31], TarBase [32], targetscan[33], and miRanda[34] predicted data.

### EXPRESSION ANALYSIS

Spearman correlation analysis was performed to determine the relationship between expression in CNV and mRNA, as well as the correlation between methylation levels and mRNA expression levels. Differential analysis was completed between the methylation of tumor and normal samples. Single gene expression differential was also analyzed. The significance difference between two group comparisons was evaluated using the t-test. mRNA expression data of the gene set was also analyzed with the Gene Set Variation Analysis (GSVA) using the Wilcoxon test and the ANOVA t-test.

### SURVIVAL ANALYSIS

SNV data and clinical survival data were separated into groups when the specific gene was mutated in the samples and was analyzed using the Cox Proportional-Hazards model. The CNV data, which was processed with GISTIC 2.0 [35], and clinical data were separated into groups based on amplification, deletion, or unaltered. Within these groups, R package survival was used to determine the survival status and time. The various groups’ survival differences were tested by using Logrank tests and the Cox proportional hazards model. Clinical survival data and methylation were analyzed. The data was divided into high and low methylation groups. The risk ratio (Hazard ratio, HR) was calculated using the Cox proportional hazards model, and a Log Rank test was performed to see whether the survival differences between the groups were statistically significant. Clinical survival data and mRNA expression data were also analyzed; tumor samples were divided into high and low-expression groups using median mRNA values. R package survival was used to fit survival time and status between groups, and the Cox Proportional-Hazards model and Logrank tests were performed for every gene in every cancer.

### PATHWAY ANALYSIS

Differences in gene expression between pathway activity groups were also studied. Scores were calculated from 7876 samples using the reverse phase protein array (RPPA) from the TCPA database. Using this information, ten cancer-related pathways were included to calculate the pathway activity score: hormone estrogen receptor (ER), hormone androgen receptor (AR), epithelial-mesenchymal transition (EMT), (TSC)/mechanistic target of rapamycin (mTOR), receptor tyrosine kinase (RTK), phosphatidylinositol-4,5-bisphosphate-3-kinase (PI3K)/protein kinase B (AKT), RAS/mitogen-activated protein kinase (MAPK), cell cycle, and apoptosis pathways. The sum of all the relative protein levels minus the negative regulatory components of a particular pathway is then the pathway score. The student T-test analyzed pathway activity scores (PAS), which were calculated in previous studies [36,37]. The P value was adjusted by the FDR; when the FDR<=0.05, it is deemed significant. Samples were divided into high and low by median gene expression. When PAS (Low expression of Gene A) < PAS (High expression of Gene A), it is regarded that the gene would have an activating effect on a pathway. If not, then it would possess an inhibitory effect on a pathway.

### MicroRNA REGULATION ANALYSIS

MiRNA-gene pairs with recorded data in databases were calculated. Gene expression and miRNA expression were merged via TCGA barcode, and the association between the two was tested by Pearson product-moment correlation coefficient and the t-distribution. Taking into account the presence of positive regulators such as transcription factors, a miRNA exhibiting a negative correlation will be regarded as a potentially negatively regulating pair. The P-value was adjusted by the FDR so that the significant genes remained. The miRNA regulation network was constructed by the visNetwork R package.

### IMMUNOHISTOCHEMISTRY EXPERIMENTAL VALIDATIONS

Immunohistochemistry data were sourced from the Human Protein Atlas (HPA) database. Different samples or cell types within a tissue were integrated by assigning scores (high=3, medium=2, low=1, not detected=0) followed by averaging. The differences in protein expression between cancerous and non-cancerous tissues, based on immunohistochemistry experiments, were compared and visualized in a heatmap. Another heatmap was created to show the prognostic association of protein expression across various cancer types. This analysis correlated expression levels (high, medium, low, not detected) with patient survival, considering a p-value of less than 0.05 as significant.

### CHEMOTHERAPEUTIC AGENTS SENSITIVITY CORRELATION ANALYSIS

We collected the IC50 of 4 commonly used cancer chemotherapeutic agents (Cisplatin, Paclitaxel, Doxorubicin, and Gemcitabine) in all cell lines and its corresponding mRNA gene expression from Genomics of Drug Sensitivity in Cancer (GDSC). The mRNA expression data and drug sensitivity data were merged. Pearson correlation analysis was performed to get the correlation between gene mRNA expression and drug IC50. P-value was adjusted by FDR.

## RESULTS

### SINGLE NUCLEOTIDE VARIATION ANALYSIS

The single nucleotide variation (SNV) profiles of the protein kinase genes associated with the mitotic DNA integrity checkpoint signaling were analyzed. The SNV profile showed a large majority of these genes have a high SNV mutation frequency across some cancer types, such as UCEC and SKCM. Specifically, PRKDC and ATM were the most frequently mutated genes. ATM was mutated in approximately 19% of all the UCEC samples. UCEC generally had the higher SNV mutation frequency compared to the cancer types, though SKCM, COAD, and STAD all showed relatively high SNV mutation frequencies for many genes. **(Figure 2A)** According to the SNV landscape, the vast majority of the SNV mutations were missense mutations. However, the second leading mutation type was nonsense mutations. (Figure 2B) The profile of the survival association with the SNV of the genes revealed that there may be a relationship between CHEK1 in BRCA as it relates to patient survival times. It also revealed that LICH and COAD were more likely to associate with the SNV of these genes. Even so, only a few genes were found to be significant, and most other cancer types found no significance. (Figure 2C) Based on the SNV analysis, while there were high SNV frequencies for many genes, it is suggested that SNV in protein kinases related to the mitotic DNA integrity checkpoint signaling might not be critical for most cancers; but, it may have some impact on BRCA.

**Figure 2:**
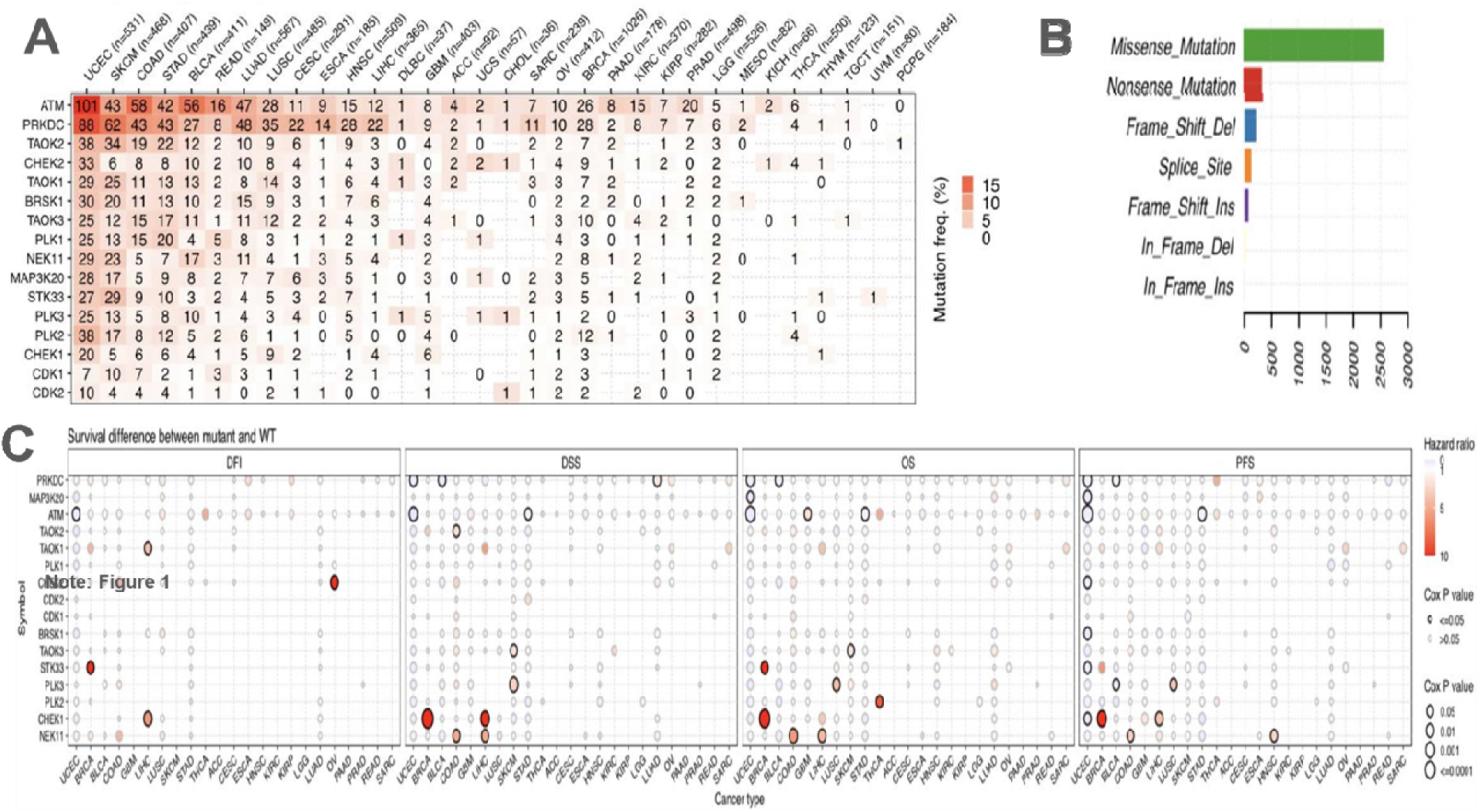
A: Heatmap of the mutation frequency. Numbers associate the number of samples containing mutation frequencies in a given cancer; 0 signifies that there are no mutation frequencies in that region of the genome. Color correlates with the frequency of mutation amount per cancer. B: Graph of mutation types of the genes in cancers. C: SNV survival plot between mutant and normal genes.

### COPY NUMBER VARIATION PROFILE ANALYSIS

The gene set’s copy number variation (CNV) profiles were analyzed. The CNV analysis revealed a great variation of the patterns in the genes across various cancer types. The CNV pie chart and plots showed that there is a large abundance of both homozygous and heterozygous CNV. However, it appears to be that there are more heterozygous CNVs than homozygous across cancer types. (Figure 3 and S-Fig.1). The pan-cancer CNV survival analysis profiles unveiled that the CNV of many of the genes is associated with patient survival. The cancer types these genes associate the most with are UCEC, KIRP, and LGG. Specifically, it was revealed that CDK2 & TOAK2 in KICH and NEK11 in KIRP might have the most significance when related to survival. (Figure 4A) The correlation profile between CNV and mRNA expression reveals that CNV is positively correlated with the majority of the genes. (Figure 4B). Based on the CNV analysis, there may be a potential association between heterozygous CNV of th protein kinase genes in the mitotic DNA integrity checkpoint signaling gene set and cancers.

**Figure 3:**
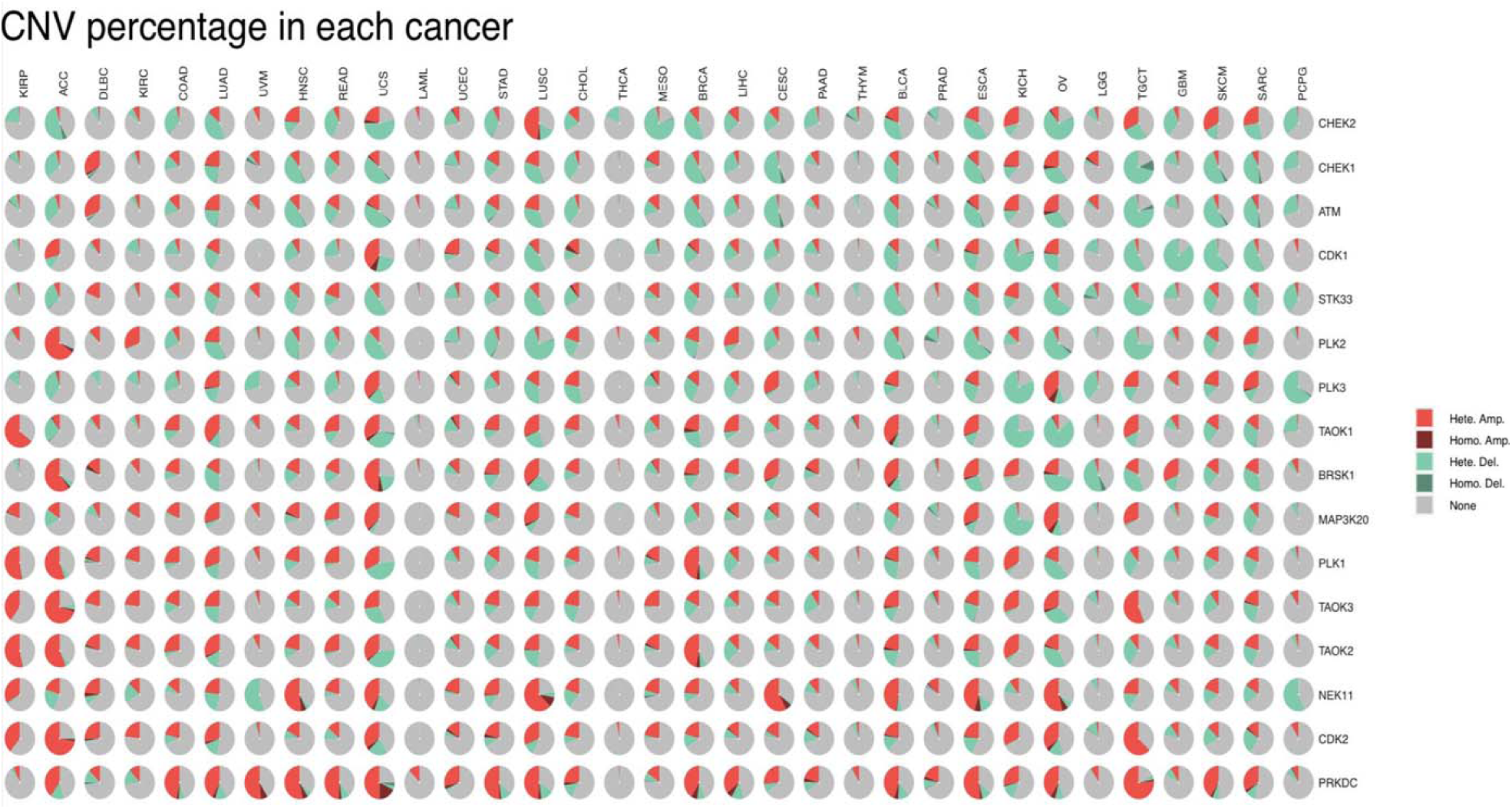
Pie charts of CNV distribution across cancers. Homo Amp = homozygous amplification; Homo Del = homozygous deletion; Hete Amp = heterozygous amplification; Hete Del = heterozygous deletion.

**Figure 4:**
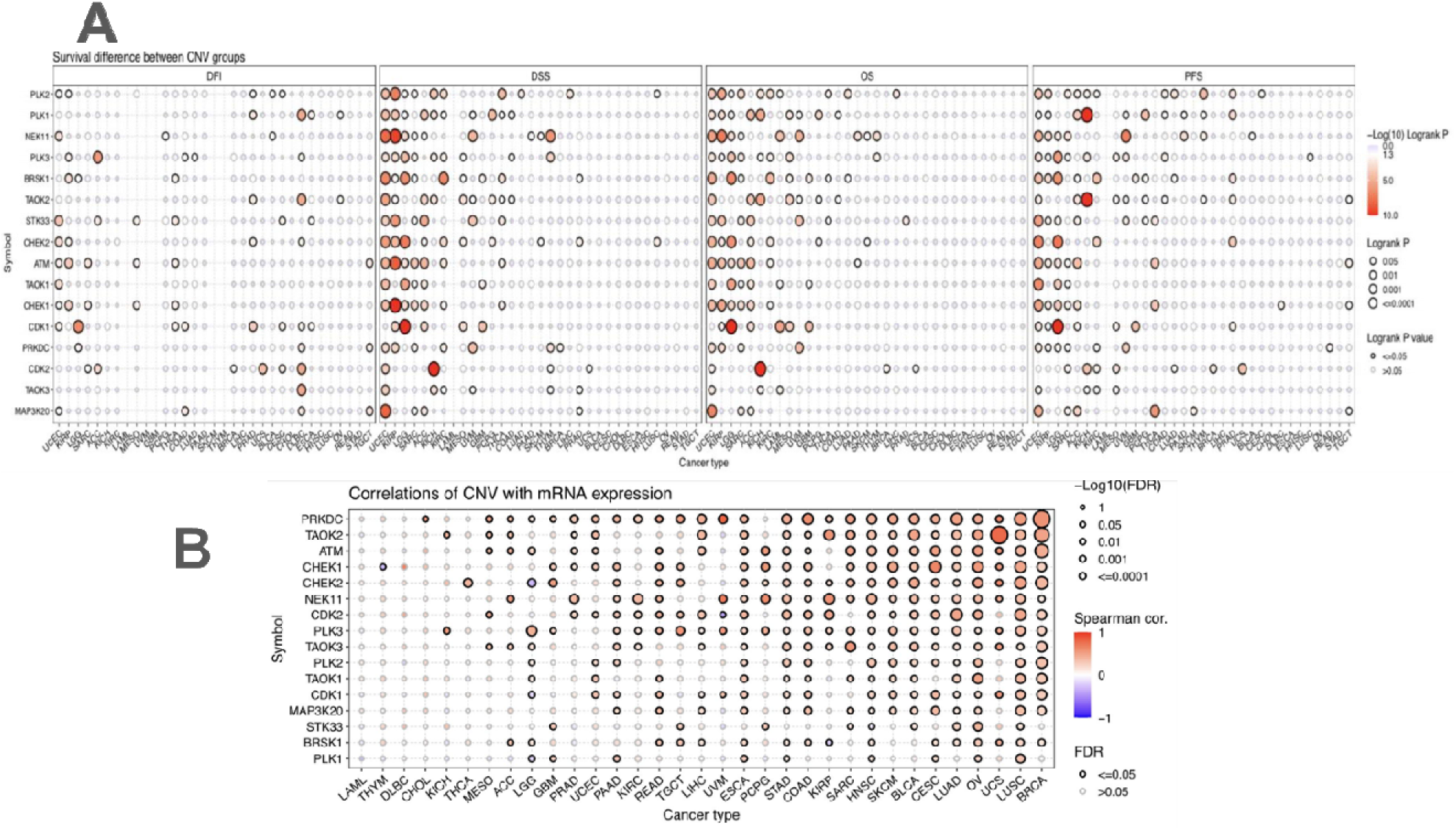
A: Survival difference between CNV groups. B: Plot of correlations between CNV and mRNA expression. Spearman cor. is the correlation coefficient.

### METHYLATION ANALYSIS

The methylation profile between tumor and normal cells revealed that some cancer cells showed a difference in methylation levels between cancer and normal tissues. STK33 gene appears to have a higher methylation level in cancer across some types than normal cells (Figure 5A). The correlation analysis between methylation and mRNA expression profiles unveiled a consistent pattern, wherein certain cancer types exhibited associations between expression and methylation. The prevailing trend in these associations was predominantly negative (Figure 5B). About the survival profile analysis, only a few cancers showed a correlation between methylation and the survival of patients (Figure 5C). The analysis suggests a potential correlation between hypermethylation and the down-regulation of genes within the specified gene set.

**Figure 5:**
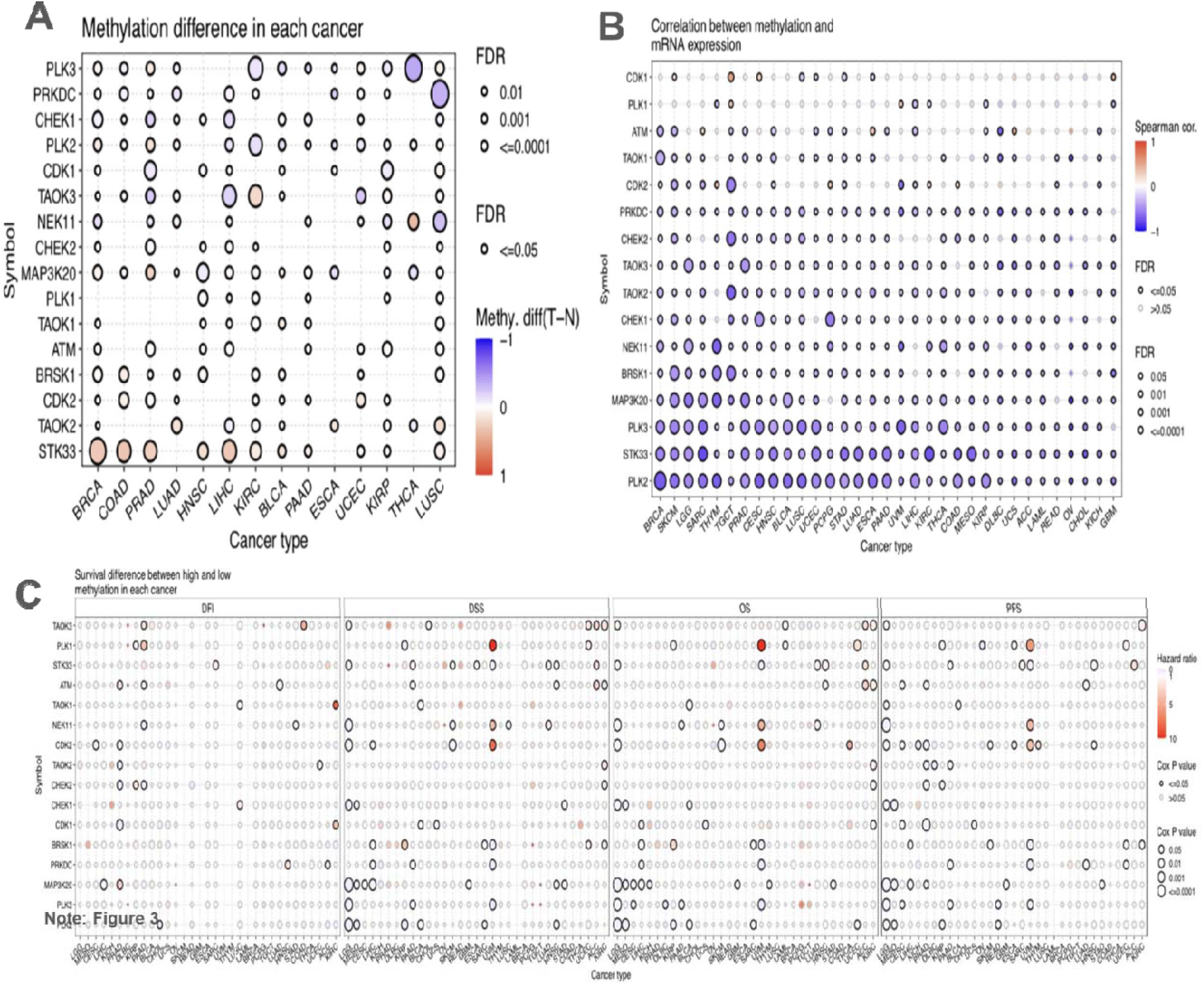
A: Methylation difference between tumor and normal tissues. B: The correlation between methylation and mRNA expression. C: Survival difference between samples with high and low methylation of the genes.

### mRNA EXPRESSION ANALYSIS

Pan-cancer expression profiles of the gene set were analyzed. We used TCGA data to examine the expression difference between cancer and non-cancer tissues of the protein kinase genes associated with mitotic DNA integrity checkpoint signaling. The findings indicated that in LUSC, PLK1, CDK1, CHEK1, and CHEK2 all appear to be up-regulated; however, PLK1 appears to be the most significant. Furthermore, PLK1 appears to be up-regulated in a few other cancer types, such as BRCA and LUAD. STK33 in KIRC and NEK11 in KICH both appear to be down-regulated, and NEK11 had a higher fold change. Despite this, the expression of these genes showed few significant differences in most cancer types when comparing cancer and non-cancer tissues (Figure 6A). The expression of these genes amongst subtypes was also analyzed. BRCA showed the most significance among the genes, but KIRC, LUAD, and STAD all appeared to have significant subtype differences as well (Figure 6B). The survival analysis showed that some genes have a relationship between expression and survival. CHEK2, CDK1, and PLK1 appear to have a larger survival risk when overexpressed in ACC. In addition, the survival analysis further shows that overexpression of PLK1 and CDK1 in KIRP may also have an association with higher survival risk of patients (Figure 6C). While there showed to be some significance across various genes, their expression levels may not be critical for some cancers.

**Figure 6:**
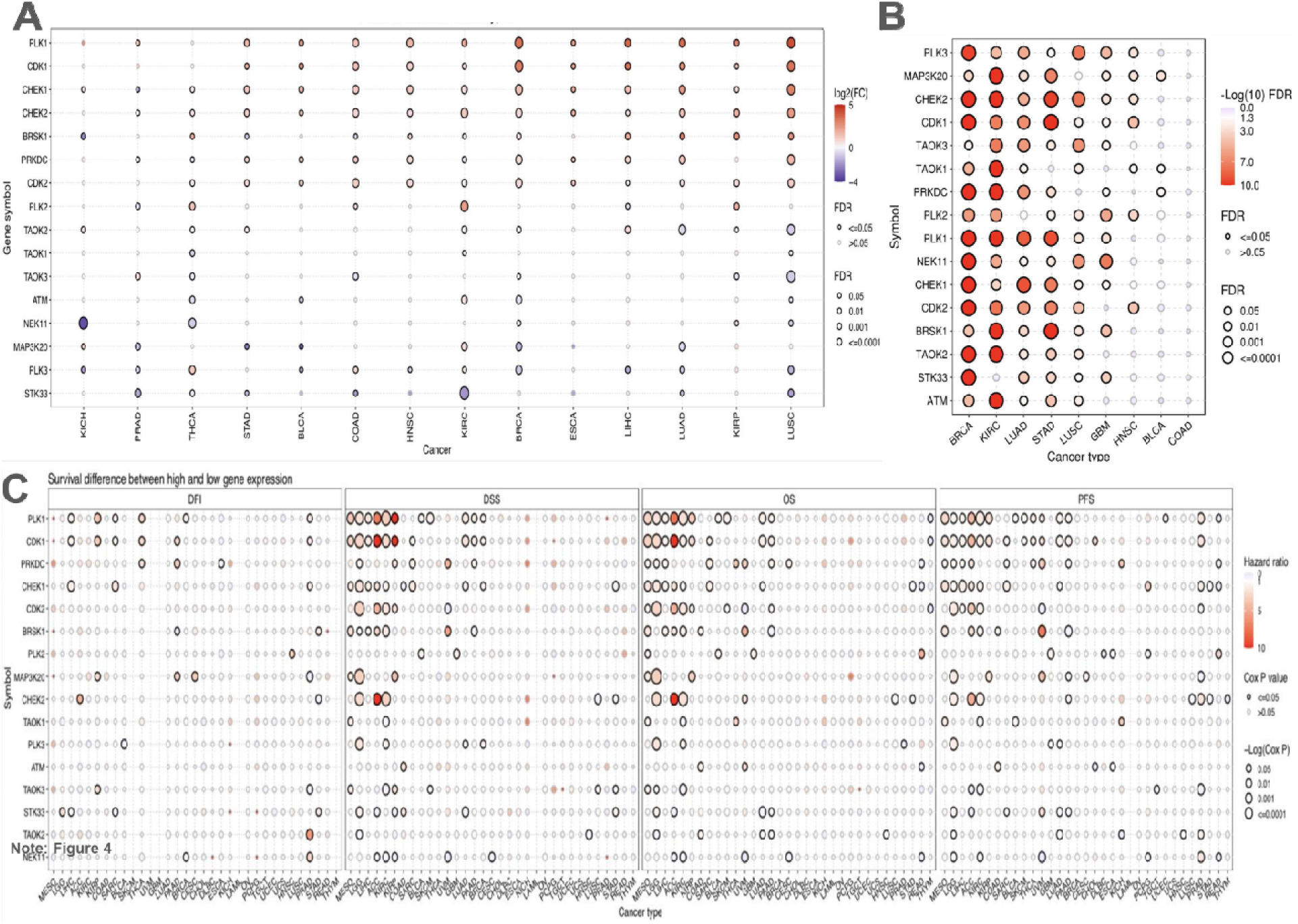
A: Expression difference between non-cancer and cancer tissues. B: Difference of expression between subtypes of cancers. C: Difference of survival between high and low gene expression.

### CROSSTALK PATHWAYS ANALYSIS

Potential crosstalk between the gene set and cancer-related pathways was analyzed. It was revealed that the mRNA expression levels of many genes had significant effects on cancer-related pathway activities. 72% of the cancer types analyzed associated high expression of CDK1 and PLK1 with cell cycle activation. Furthermore, of the cancer types analyzed, 53% associated CDK2 and 66% associated CHEK1 with cell cycle activation also. TAOK3, PLK3, PLK2, and NEK11 appear to be associated with cell cycle inhibition. PLK3 and PLK2 also seem to be associated with the inhibiting of the hormone androgen receptor. CHEK2, CHEK1, CDK2, and CDK1 appeared to relate to the inhibition of RASMAPK across cancer types (Figure 7). The pathway profiles revealed the potential relationship between protein kinase genes associated with the mitotic DNA integrity checkpoint signaling and other common cancer-related pathways.

**Figure 7:**
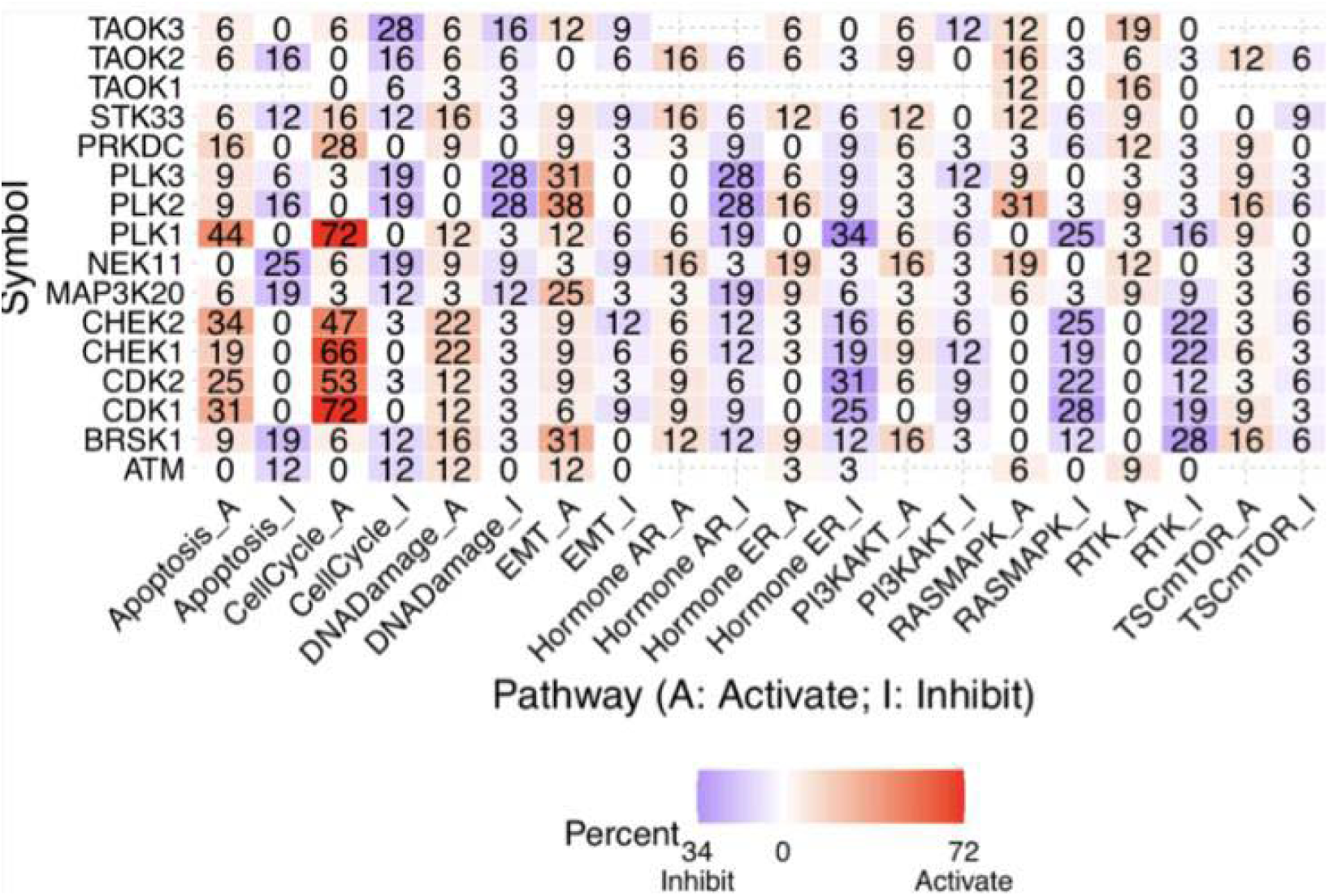
Expression and pathway activity of the genes.

### MicroRNA REGULATION ANALYSIS

MiRNA expression data was also analyzed as it relates to the protein kinases associated with the mitotic DNA integrity checkpoint signaling gene set; the data was acquired from TCGA and then assembled into a miRNA-regulation network. 15 out of 16 of the genes from this gene set were identified in the databases with regulating miRNA. It appears that multiple miRNAs, including several co-regulators, may be involved in regulating the expression of these genes. 26 miRNA regulate TAOK3, where the top regulating miRNA were hsa-miR-214-3p, hsa-miR-20a-5p, hsa-miR-93-5p, hsa-miR-106b-5p, and hsa-miR-17-5p. Has-miR-214-3p also regulates ATM; hsa-miR-20a-5p, hsa-miR-93-5p, hsa-miR-106b-5p, and hsa-miR-17-5p all regulate TAOK2 and STK33 as well. TAOK2 is regulated by 16 miRNAs; 20 total miRNA regulate STK33, where hsa-miR-994 appears to be a top regulator. Hsa-miR-994 cross-regulates with ATM as well, also functioning as a key regulator for ATM. 18 miRNA serve as regulators for ATM. Besides hsa-miR-994, other major miRNA regulators for ATM were hsa-miR-141-3p, hsa-miR-18a-5p, hsa-mir-194-5p, and hsa-miR-200a-3p. 17 miRNA regulate NEK11, especially by hsa-miR-24-3p, hsa-miR-590-5p, and hsa-miR-3140-3p. One of NEK11’s miRNA regulators, hsa-miR-3163, also regulates TAOK1, BRSK1, and TAOK3. 5 genes regulate BRSK1, with hsa-miR-27a and hsa-miR-27b-3p as top regulators. CDK2 appears to have 11 miRNA regulators, the most prominent being hsa-miR-200c-3p; it also cross-regulates with TAOK3 and PLK2. PLK2 is regulated by 12 miRNA and is cross-regulated with CDK2 and TAOK3 by hsa-miR-429, hsa-miR-200b-3p, and hsa-miR-200c-3p. 12 miRNA regulate CDK1, mainly hsa-miR-7-5p and hsa-miR-143-3p. The latter also regulates TAOK2. PRKDC is regulated by 7 miRNA, with hsa-miR-101-5p being its top regulator. PLK3 is regulated by 3 miRNA, while CHEK2 and PLK1 are regulated by one. CHEK1 is regulated by a total of 5 miRNA, particularly by hsa-miR-195-5p and hsa-miR-497-5p. 20 miRNA regulate TAOK1; top-regulators are hsa-miR-30e-5p, hsa-miR-30b-5p, hsa-miR-27b-3p (Figure 8). The network offered insight into miRNA potentially regulating protein kinases in mitotic DNA integrity checkpoint signaling for future referencing.

**Figure 8:**
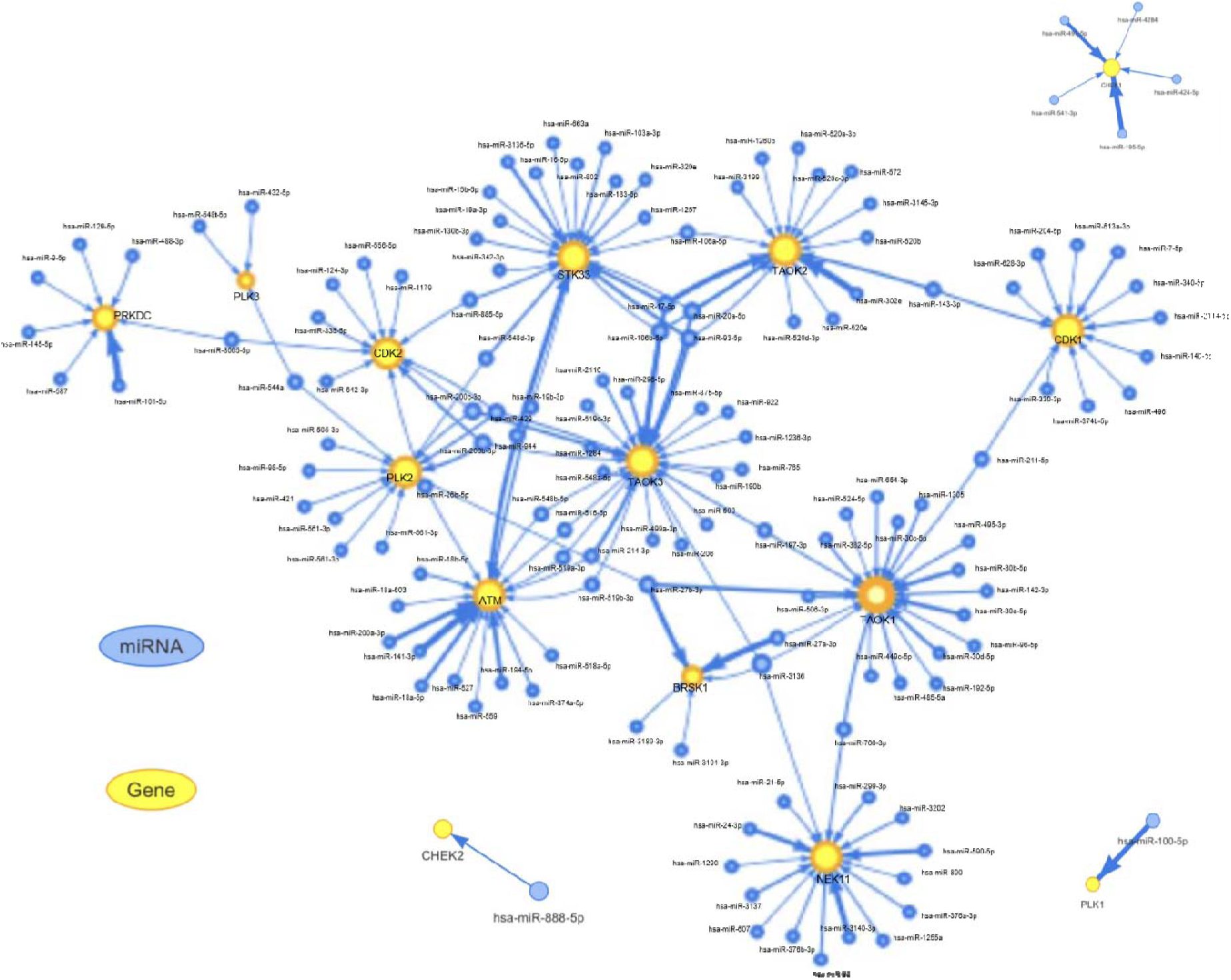
MicroRNA (miRNA) regulation network of the genes. The yellow circles represent the genes from the gene set, the blue circles represent the miRNA. The width of the node is positively correlated with its level of regulation for its associated gene.

### IMMUNOHISTOCHEMISTRY EXPERIMENTAL VALIDATIONS

We believe that an immunohistochemistry experiment to validate protein-level expression will be instrumental in verifying our conclusions. Given that our study involves a pan-cancer, multi-gene analysis, conducting experiments for all cancer types and the 16 genes in this study presents significant challenges. Fortunately, the Human Protein Atlas team has previously conducted such experiments, validating the expression of pan-cancer genes through immunohistochemistry. To further validate our conclusions, we have leveraged their prior work by integrating their experimental results with our analysis. We conducted two separate analyses: one for diagnosis, comparing protein expression in cancerous versus non-cancerous tissues, and another for prognosis, examining the biomarkers’ risky or protective properties based on survival data. This approach allows us to provide additional, relevant experimental evidence to support some of our conclusions. For cancer-non-cancer comparison, the integrated data is based on average protein expression of different patients or cell types in one sample, hence, it is not possible to calculate the significance, so this is just for general referencing. (Figure 9A) For prognosis analysis of protein expression, one of the most striking findings was that many of the protein kinases in mitotic DNA integrity checkpoint signaling are significantly risky for renal cancer. (Figure 9B)

**Figure 9:**
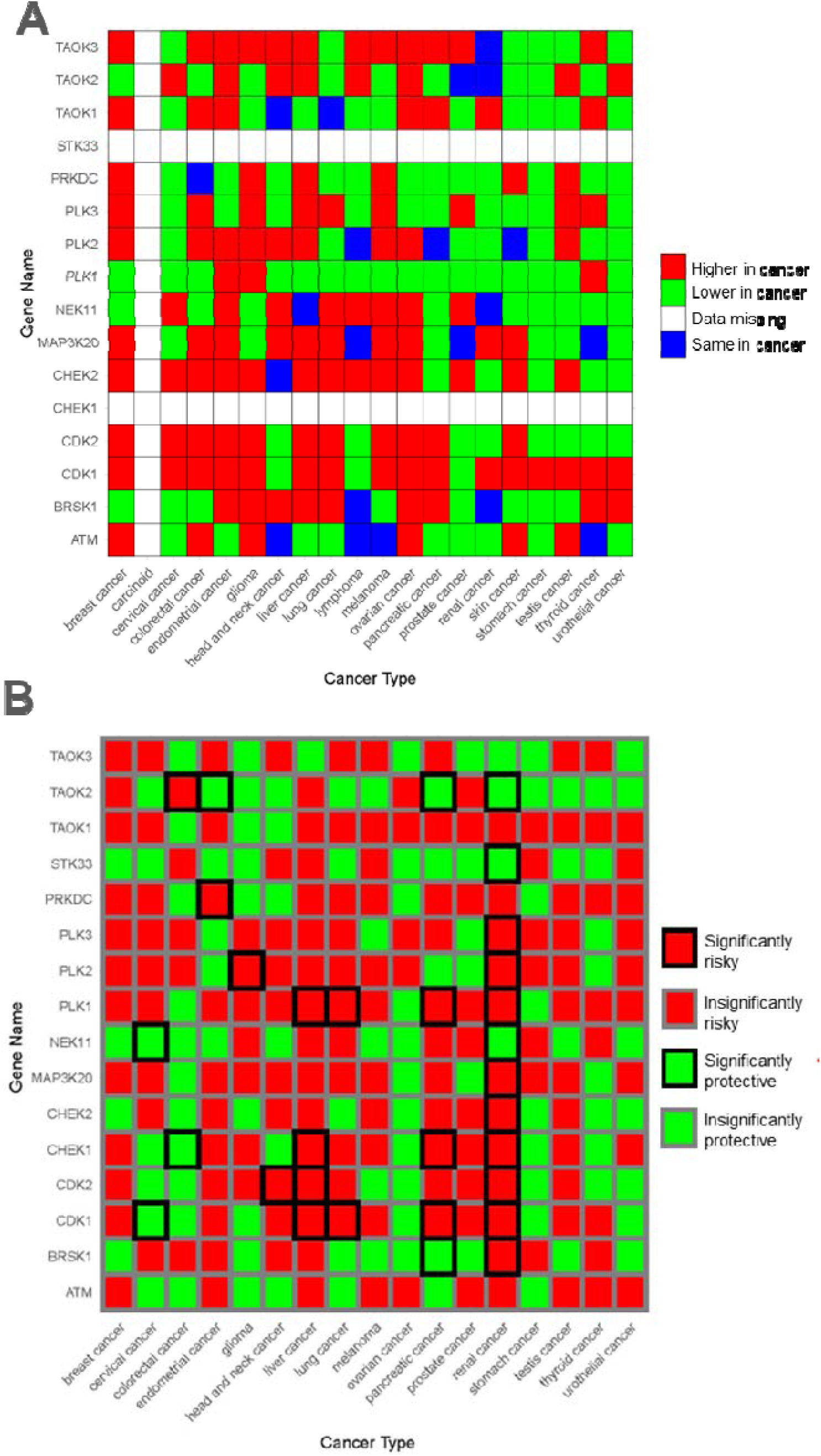
Immunohistochemistry experimental validations of the genes. Immunohistochemistry data are accessed from the Human Protein Atlas (HPA) database. A. A heatmap showing protein expression difference between cancer and non-cancer tissue based on the Immunohistochemistry experiments. B. A heatmap showing protein expression prognostic association across cancer types based on the Immunohistochemistry experiments.

### CHEMOTHERAPEUTIC AGENTS SENSITIVITY CORRELATION ANALYSIS

Understanding whether mitotic DNA integrity checkpoint protein kinases correlate with sensitivity to chemotherapeutic agents such as cisplatin is crucial for comprehending this pathway’s role in cancer treatment and drug resistance. To explore this, we utilized cell lines to evaluate gene expression and sensitivity to chemotherapy. Our study included four commonly used cancer chemotherapeutic agents: Cisplatin, Paclitaxel, Doxorubicin, and Gemcitabine. We downloaded IC50 values for these agents across all cell lines, along with corresponding mRNA gene expression data from the Genomics of Drug Sensitivity in Cancer (GDSC) database. We merged the mRNA expression data with the drug sensitivity data prior to conducting Pearson correlation analysis. The results were visualized in a volcano plot, highlighting the correlations and their false discovery rates (FDR). Our findings indicate significant correlations: TAOK2 is positively correlated with Cisplatin IC50, suggesting higher resistance, while ATM is negatively correlated with Gemcitabine, indicating increased sensitivity. Additionally, BRSK1 shows a significant positive correlation with Paclitaxel sensitivity. (Fig.10)

**Figure 10:**
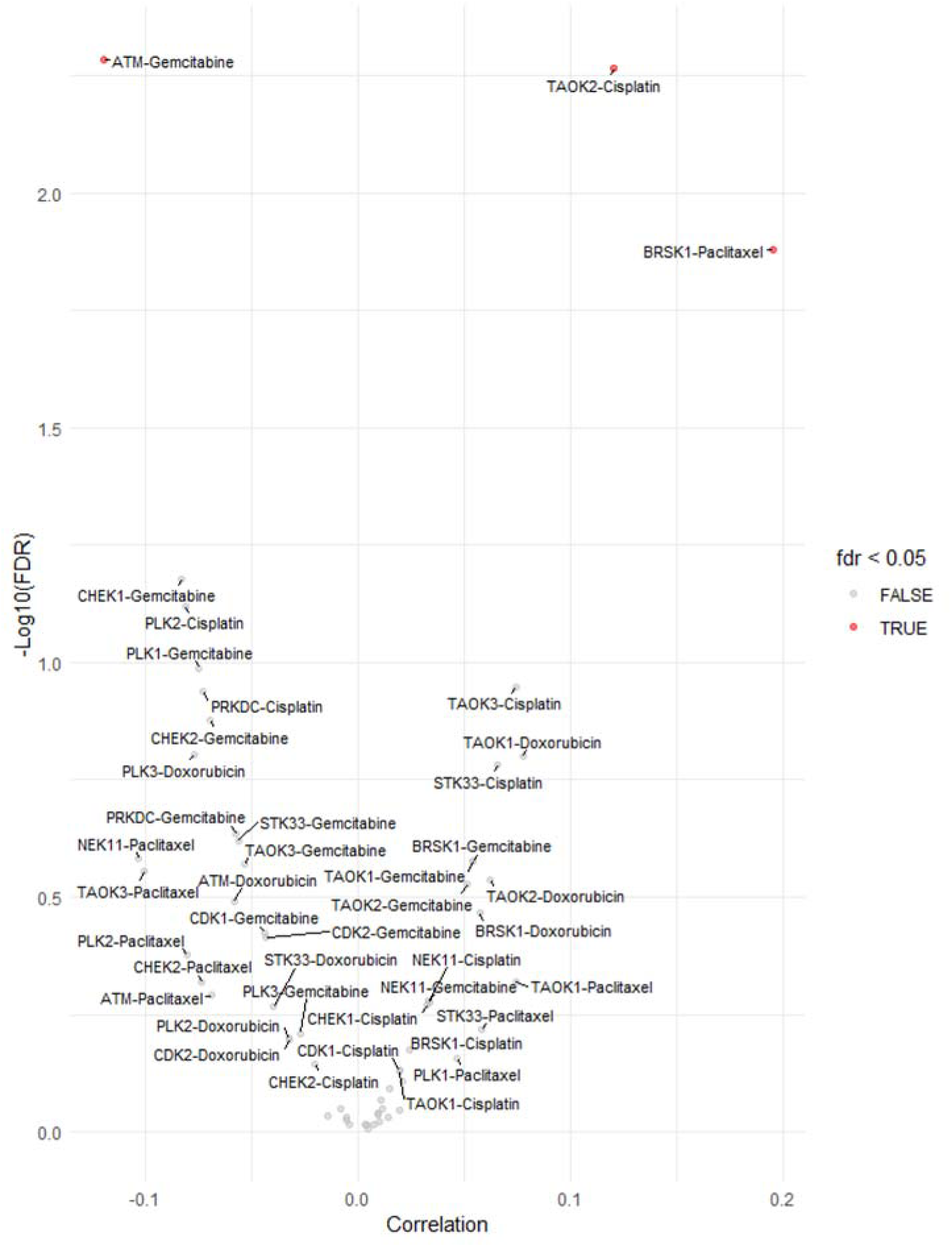
Chemotherapeutic agents sensitivity correlation analysis.

## DISCUSSION

In this study, the pan-cancer analysis of the protein kinases associated with mitotic DNA integrity checkpoint signaling showed its potential value as cancer prognostic and diagnostic biomarkers. Our survival results regarding SNV displayed that CHEK1 has an impact on the survival time of BRCA patients. Previous studies [38] have also found that CHEK1 is upregulated in BRCA tissues compared to normal breast tissues, and patients with high levels of CHEK1 mRNA or CHEK1 alterations are more likely to receive a worse prognosis. CHEK1 could be a potential biomarker for BRCA, as our results indicated it to be very significant and risky. However, our mutation frequency heat map revealed that only a very small portion of patients had expressed the gene, so we suggest future studies to pursue further research on this front.

Our correlation results between mRNA expression and methylation showed that hypermethylation negatively affects mRNA expression. These outcomes were similar to previous studies that indicated hypermethylation is associated with repressed transpiration [39]. This can limit the production of proteins in the cell, including kinases, which means the necessary signals the kinases perform to maintain DNA integrity will not be executed. This could result in aneuploidy in cells, for DNA could pass through this portion of the cell cycle unchecked.

In this study, it was revealed that there is a correlation between the high expression of some mitotic DNA integrity checkpoint signaling genes and the survival time of patients. PLK1 and CDK1 were shown to have a larger survival risk when overexpressed in KIRP. Previous studies have agreed that these genes have an association with tumorigenesis of KIRP cells [40,41], and have a potential use as a target in cancer therapies.

Our study reveals that certain protein kinases in the mitotic DNA integrity checkpoint signaling pathway influence well-established cancer pathways. Notably, CDK1 was observed to activate the cell cycle pathway in 72% of the sampled cancer types. This observation aligns with findings from prior studies, as CDK1 phosphorylates downstream substrates that play a significant role in cancer progression [42]. We propose that future research delve deeper into this connection.

While most of the mRNA expression appeared to correlate with methylation, CDK1 did not show much significance. However, in the miRNA-gene pair analysis, CDK1 was shown to be regulated by 12 miRNA. We think miRNA pathways potentially have an impact on the regulation of CDK1 in cancers. Previous studies have concurred with the notion that miRNA plays a role in regulating CDK1 [43], as the reason might be that miRNA binds to certain parts of the mRNA that codes for CDK1, leading to repressed transcription of the protein kinase.

There may be some possible limitations to this study. For instance, only tissue samples were analyzed. While they show valuable snapshots of the cancer, they do not capture the evolving state of the disease. Previous studies have revealed relationships among genes, diseases, and environmental factors such as microbiotics or drug treatments[44–66]. However, whether the role of protein kinases associated with mitotic DNA integrity checkpoint signaling can be influenced by these factors requires further exploration. This understanding is crucial for developing targeted therapies and improving existing treatment strategies, particularly in the context of personalized medicine. Exploring these interactions may provide new insights into how these kinases can be modulated by external influences, potentially leading to breakthroughs in cancer treatment and drug resistance management. The results of this study provide valuable information, as they offer useful insights for potential diagnostic and prognostic biomarkers, as well as potential targeted therapies in a clinical setting. These profiles also give an overall picture of the protein kinases in the mitotic DNA integrity checkpoint signaling gene set for future references in cancer research.

## CONCLUSION

This study provided insight into SNV, CNV, methylation, mRNA expression, pathway crosstalk, and miRNA regulation profiles for the protein kinases associated with the mitotic DNA integrity checkpoint signaling across 33 different cancer types. The profiles can provide an understanding of future cancer treatment as potential prognostic and diagnostic biomarkers, such as CHEK1 in BRCA. It also revealed gene-miRNA relationships and connections between other cancer-related pathways. The protein kinases in mitotic DNA integrity checkpoint signaling may play a role in cancer development and, with adequate research, could potentially be developed as biomarkers for cancer diagnosis and prognosis. However, further efforts are necessary to validate their clinical value for diagnosis and prognosis and to develop practical applications in clinical settings. Nevertheless, these pan-cancer profiles offer a better overall understanding as well as useful information for future reference regarding mitotic DNA integrity checkpoint signaling in cancer.

## Declarations

### Author contributions

CONCEPTION: Hengrui Liu

INTERPRETATION OR ANALYSIS OF DATA: Ayana Meegol Rasteh and Hengrui Liu

PREPARATION OF THE MANUSCRIPT: Ayana Meegol Rasteh

REVISION FOR IMPORTANT INTELLECTUAL CONTENT: Hengrui Liu and Panpan Wang

SUPERVISION: Hengrui Liu and Panpan Wang

### Availability of data and materials

The source of the raw data was provided in the paper and the raw analysis data of this study are provided by the corresponding author with a reasonable request.

### Ethical approval

Not applicable.

## Acknowledgments

We thank Melody Fallah-Khair, Farzin Rasteh, Weifen Chen, Zongxiong Liu, Bryan Liu, and Yaqi Yang for their support and contribution.

## List of the cancer-type abbreviations

ACC: Adrenocortical carcinoma
BLCA: Bladder Urothelial Carcinoma
BRCA: Breast invasive carcinoma
CESC: Cervical squamous cell carcinoma and endocervical adenocarcinoma
CHOL: Cholangio carcinoma
COAD: Colon adenocarcinoma
DLBC: Lymphoid Neoplasm Diffuse Large B-cell Lymphoma
ESCA: Esophageal carcinoma
GBM: Glioblastoma multiforme
HNSC: Head and Neck squamous cell carcinoma
KICH: Kidney Chromophobe
KIRC: Kidney renal clear cell carcinoma
KIRP: Kidney renal papillary cell carcinoma
LAML: Acute Myeloid Leukemia
LGG: Brain Lower Grade Glioma
LIHC: Liver hepatocellular carcinoma
LUAD: Lung adenocarcinoma
LUSC: Lung squamous cell carcinoma
MESO: Mesothelioma
OV: Ovarian serous cystadenocarcinoma
PAAD: Pancreatic adenocarcinoma
PCPG: Pheochromocytoma and Paraganglioma
PRAD: Prostate adenocarcinoma
READ: Rectum adenocarcinoma
SARC: Sarcoma
SKCM: Skin Cutaneous Melanoma
STAD: Stomach adenocarcinoma
TGCT: Testicular Germ Cell Tumors
THCA: Thyroid carcinoma
THYM: Thymoma
UCEC: Uterine Corpus Endometrial Carcinoma
UCS: Uterine Carcinosarcoma
UVM: Uveal Melanoma

